# Minute-scale Periodicity of Neuronal Firing in the Human Entorhinal Cortex

**DOI:** 10.1101/2022.05.05.490703

**Authors:** Zahra M. Aghajan, Gabriel Kreiman, Itzhak Fried

## Abstract

Grid cells in the entorhinal cortex demonstrate spatially periodic firing patterns, which are thought to provide a map of space on behaviorally relevant length scales. It is unknown, however, whether such periodicity exists for behaviorally relevant time scales in the human brain. Here we investigated neuronal firing during a temporally continuous uninterrupted experience by presenting fourteen neurosurgical patients with an audiovisual video while recording single neuron activity from multiple brain regions. We report on a set of units that modulate their activity in a strikingly periodic manner across different timescales—from seconds to many minutes. These cells were most prevalent in the entorhinal cortex. Time within the video could be decoded from their population activity. Furthermore, these cells remapped their dominant periodicity to shorter timescales during a subsequent recognition memory task. When the audiovisual sequence was presented at two different speeds (regular and faster), a significant percentage of these temporally periodic cells (TPCs) maintained their timescales, suggesting a degree of invariance with respect to the narrative content. The temporal periodicity of TPCs may complement the spatial periodicity of grid cells Whether these cells provide scalable spatiotemporal metrics for encoding and retrieval of human experience warrants future investigations.

## Introduction

Integrating the content of human experience in space and time constitutes the basis for our remarkable ability for episodic memory and mental time travel(*1–4*). In rodents, several temporal coding schemes involving the hippocampal-entorhinal circuitry have been reported(*5–14*), including: (a) “time cells” in the hippocampus and medial entorhinal cortex (MEC) firing at specific points in time during a short timed interval(*5–7*); (b) “ramping cells” in the lateral entorhinal cortex (LEC) whose ramping firing activity enables extraction of time for distinct experiences during the task(*14*); (c) “event-specific” cells in the hippocampus coding for temporal order of events(*12*); and (d) degradation in the population of place cells’ activity over hours and days (*8–11*). Together, the firing properties of these cells—i.e., their sequential activation or their activity decay at different timescales—with respect to experimental temporal boundaries, are thought to provide timestamps of episodic memory.

Considering the temporal representation in the human hippocampal-entorhinal system, time can be regarded as an additional dimension to space. Grid cells in the entorhinal cortex provide a scalable map with spatial periodicity(*15, 16*) when animals forage freely for food in an open environment. To reveal an analogous temporal periodicity would require more naturalistic scenarios where time is studied at multiple timescales over prolonged periods spanning seconds to many minutes. Many perception and episodic memory experiments are dominated by a controlled stimulus-response methodology—requiring intermittent sensory input and subject response—therefore, disrupting the natural temporal continuity of behavior.. If such temporal periodicity existed, one would expect that spatial grid properties—such as rate remapping with environmental changes, and distinct grid modules with different spatial scales—would translate into the time domain. Indeed, this hypothesis is consistent with recent accounts on the role of rodent MEC in interval timing and the idea of “navigating through time” (*17–19*).

Although temporal periodicity has been observed in many aspects of biological systems, for example cardio-respiratory signals in the seconds scale and neural oscillations in the subseconds range (e.g., theta, beta, gamma oscillations), the presence of neural representations on longer timescales deserves investigation. Here we sought to investigate the existence of temporal periodicity in timescales that are relevant for human experience and behavior. We created a realistic immersive flow of information along extended temporal scales—by employing a paradigm with uninterrupted audiovisual sequence—while we recorded units’ activity in multiple brain regions in humans.

## Results

### Behavioral task

Participants were fourteen neurosurgical patients (age = 31±9; 9 Female) with intractable epilepsy who were implanted with intracranial depth electrodes in order to identify the seizure focus for potential subsequent surgical cure. First, we recorded spiking activity from microwires while nine of the fourteen participants watched a 42-minute movie (first episode, season six of “24” TV series; Fig. 1a)(*20*) and performed a recognition memory test afterwards. During the memory test, they were presented with brief movie shots and were asked whether they had seen the clip before. The target movie shots were randomly interleaved with an equal number of foil movie shots (chosen from the second episode of “24” that the patient had not seen, Fig. 1a, b; for further detail see Methods, Behavioral Tasks).

**Figure 1:**
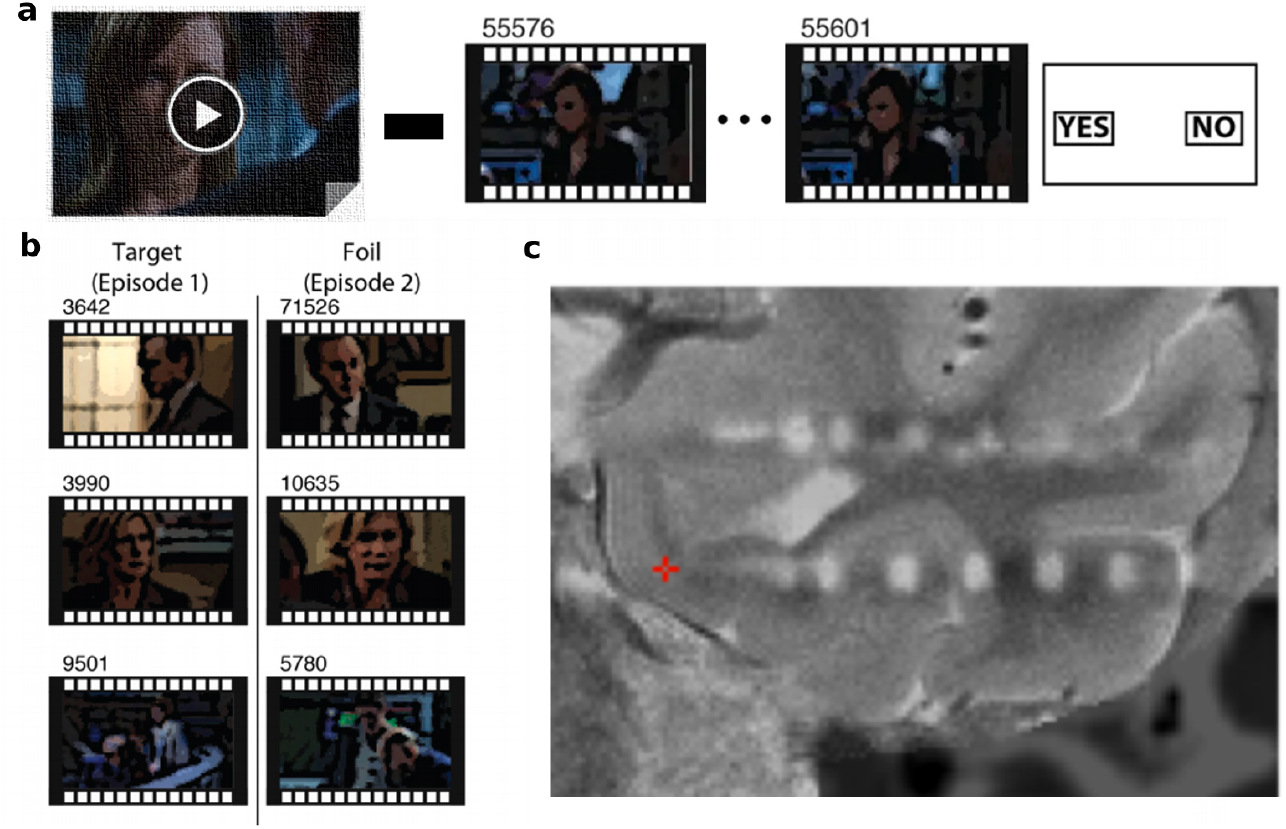
Task structure and electro-physiological recordings. **a**. Participants watched an episode of the “24” TV series (∼42 minutes in duration) and afterwards they were tested for recognition memory where they were shown short clips and asked whether they had previously seen them. **b**. The memory test included target clips (taken from the same episode they had watched, left column) and foil clips (taken from the next episode they had never seen before, right column). Images are adapted and modified from a previous publication(*20*). **c**. Depth electrodes were localized by co-registering high resolution post-operative CT images with high resolution preoperative MRI. Red crosshair indicates the location of a microwire in the left entorhinal cortex.

### Units showed periodic modulation of firing in time

We identified 382 units with a minimum firing rate of 0.05Hz (mean rate = 1.55, [0.46, 3.67] Hz) using previously described methods(*21–24*) (Methods, Data Acquisition). In order to localize these units for each participant, a high-resolution post-operative CT image was co-registered to a pre-operative whole brain and high-resolution MRI and the location of the microwires were determined for each depth electrode (Fig. 1c; Methods, Electrode Localization). These units were thus localized to eleven unique regions (Table S1) with almost half of the units (49.60%) recorded from medial temporal lobe (MTL) regions (Table S2). To display the firing rate of each unit, we binned the spikes into 100ms segments and applied a Gaussian smoothing kernel with 500ms width, followed by division by the duration of the time bin (Fig. 2a; Methods, Electrophysiological Analyses). Visual examination of the firing rates revealed that some units exhibited striking periodicity in their firing over the course of the movie, and the timescale of this periodicity varied from unit to unit, ranging from tens of seconds to several minutes (Fig. 2a). This periodicity was further demonstrated by the peaks observed in the autocorrelogram of each unit’s firing rate in time (Fig. 2b; Methods, Electrophysiological Analyses). Additionally, we used Generalized Linear Models (GLMs) to capture the time-varying firing rate as a Poisson process using basis functions that were periodic in time and inspected the model fit as well as the basis functions that were significant in explaining the rate (Fig. 2a, Fig. S1; Methods, Electrophysiological Analyses). The firing rate of these cells oscillated with a periodicity centered around one or more characteristic frequencies. We refer to these cells as Temporally Periodic Cells (TPC) given that their firing rate appears periodic in time.

**Figure 2:**
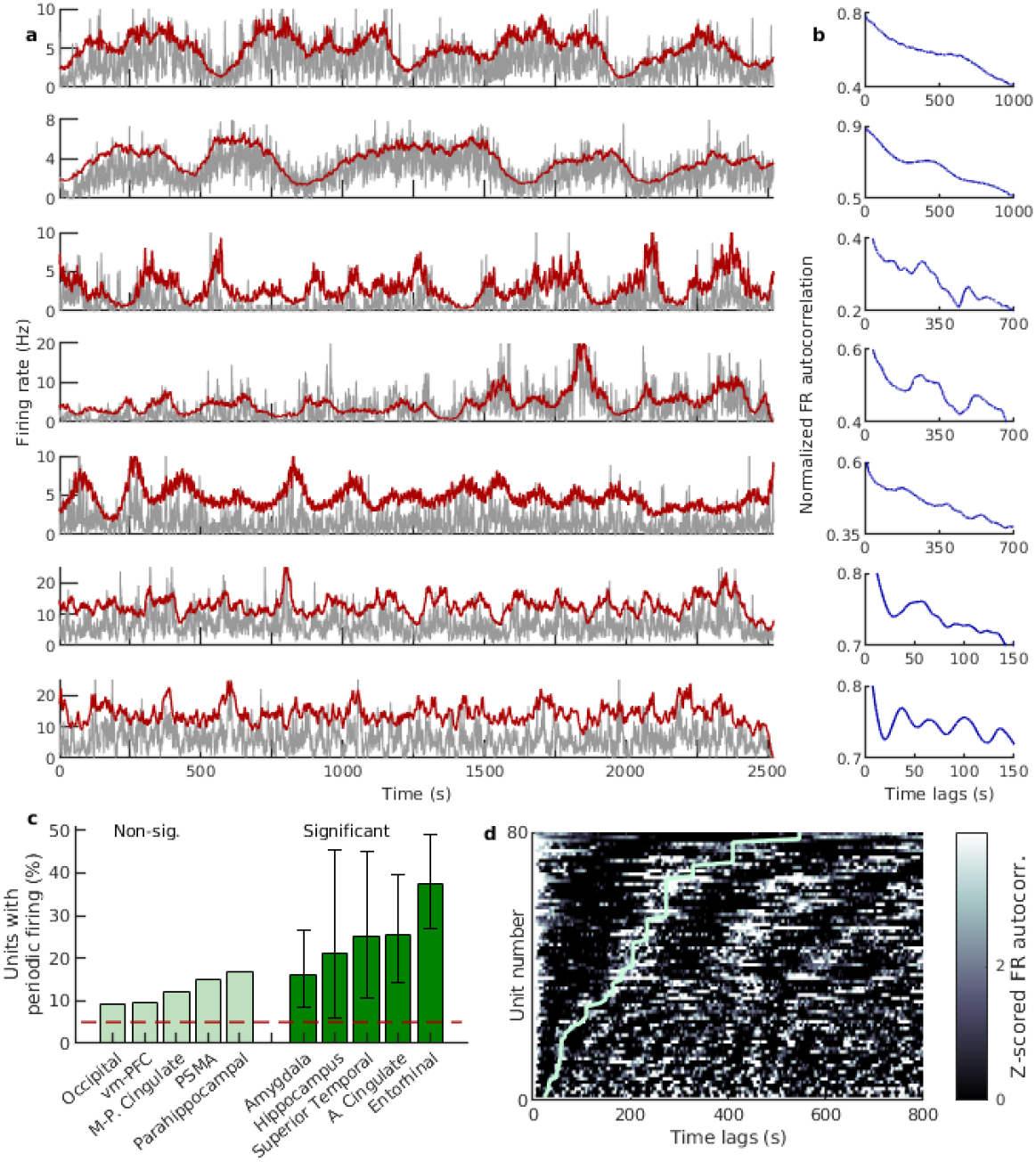
Temporally Periodic Cells (TPCs) exhibited significant periodic firing during movie viewing. **a**. Seven example TPCs firing activity. These units were recorded from ventromedial prefrontal cortex (vm-PFC), entorhinal cortex (EC), EC, anterior cingulate, EC, EC, and parahippocampal gyrus respectively. Gray line indicates the firing rate (smoothed spike train divided by the 100ms time bin). Red line indicates the GLM fitted firing rate (see Methods). **b**. Each row is the normalized autocorrelation of the smoothed firing rate of the unit shown in (a). Note the local peaks in the autocorrelograms (showing a periodicity in the unit firing), as well as the different timescales for each unit (x-axis limits are adjusted according to the unit’s timescale). The autocorrelations are smoothed only for visualization purposes. **c**. Within each recording region, the percentages of TPCs during movie viewing are shown in green bars and the error bars indicate the confidence intervals of a binomial test (for a full list of the number of recorded units and significant TPCs per region, see Table S2). The EC region had the largest percentage of TPCs and the regions marked in light green did not have a significant percentage of TPCs (the confidence intervals of the binomial test included the 5% chance level). The percentage of TPCs in regions other than EC are not within the confidence intervals of the EC region. **d**. Z-scored autocorrelations of all the TPCs’ firing rates (colormap; N = 80) were sorted by the dominant periodicity (light green line, see Methods) for each unit (each row). Note the visible diverging lines parallel to the dominant period, corresponding to periodicity in the signal. The dominant periodicity of the units shown in (a) are as follows: 546.14s, 409.60s, 273.10s, 273.10s, 182.04s, 56.50s, 34.86s.

To quantitatively assess the extent to which neurons fired in a periodic fashion, we computed the autocorrelation of the firing rate for each unit and compared it against the null hypothesis constructed using shuffled data (specifically, the autocorrelations computed over the shuffled firing rates of the same unit; for details see Methods, Electrophysiological Analyses). A unit with an autocorrelation value outside the [2.5%, 97.5%] of the shuffled data was identified as a putative TPC. Furthermore, we used a cluster-based permutation test to correct for multiple comparisons in identifying these units and found a total of 80 TPCs (Fig. S2a; more examples of TPCs are shown in Fig. S2, S3). We then quantified the percentage of TPCs within each region and found that multiple regions contained a significant fraction of these units, with the entorhinal cortex holding the largest population of TPCs (30 out of 80 total entorhinal units; 37.50%, [26.92, 49.04]%, confidence intervals from binomial test), followed by the anterior cingulate region (13 out of 51 total units; 25.49%, [14.33, 39.63]%, confidence intervals from binomial test)(Fig. 2c, Table S2).

### TPCs’ periodicities span multiple timescales

We next asked how the periodicity of TPCs varied across units. We calculated the dominant period for each unit as follows: 1) the firing rate autocorrelation was z-scored with respect to the shuffled data for each unit; 2) an FFT was performed on the z-scored autocorrelation values; and 3) the period at which the largest power was contained was determined to be the dominant period for that unit (Fig. S2; Methods, Electrophysiological Analyses). This approach allowed us to examine the periodicity scale of all TPCs. The population activity of these units spanned temporal scales ranging from tens of seconds to several minutes (Fig. 2d). It is worth noting that such large temporal scales are beyond the temporal response patterns of traditional time cells, previously observed in the hippocampus and medial entorhinal cortex (MEC) of rodents, which involved temporal scales on the order of a few seconds(*6, 7*). The timescales of TPCs are more similar to those of the ramping time cells discovered in the rodent LEC(*14*).

The population of TPCs exhibited multiple timescales even within each participant (Fig. 3a; Table S3), as well as within different regions (Fig. 3b). At the population level, the distribution of the TPCs’ dominant periods revealed a non-uniform distribution (p < 10^−5^; single sample Kolmogorov-Smirnov test against uniform distribution) and some timescales appeared to be more pronounced (e.g., dominant periodicities at 62.5s, 112.5s, 180s, 290s and 400s; Fig. 3c). Although thus far the results focused on the dominant periodicity (the oscillation with the highest power), some units had periodic firing at additional temporal scales. To determine other prominent oscillations, we calculated the relative power of the z-scored firing rate autocorrelation with respect to the power at the dominant period and found the peaks with at least 75% of the maximum power (Fig. 3d). Indeed, 35% of the units showed periodic firing at one or more frequencies in addition to their dominant periodicity (Fig. 3e). These additional frequencies were not simply multiples of each other. Few units had more than two additional frequencies.

**Figure 3:**
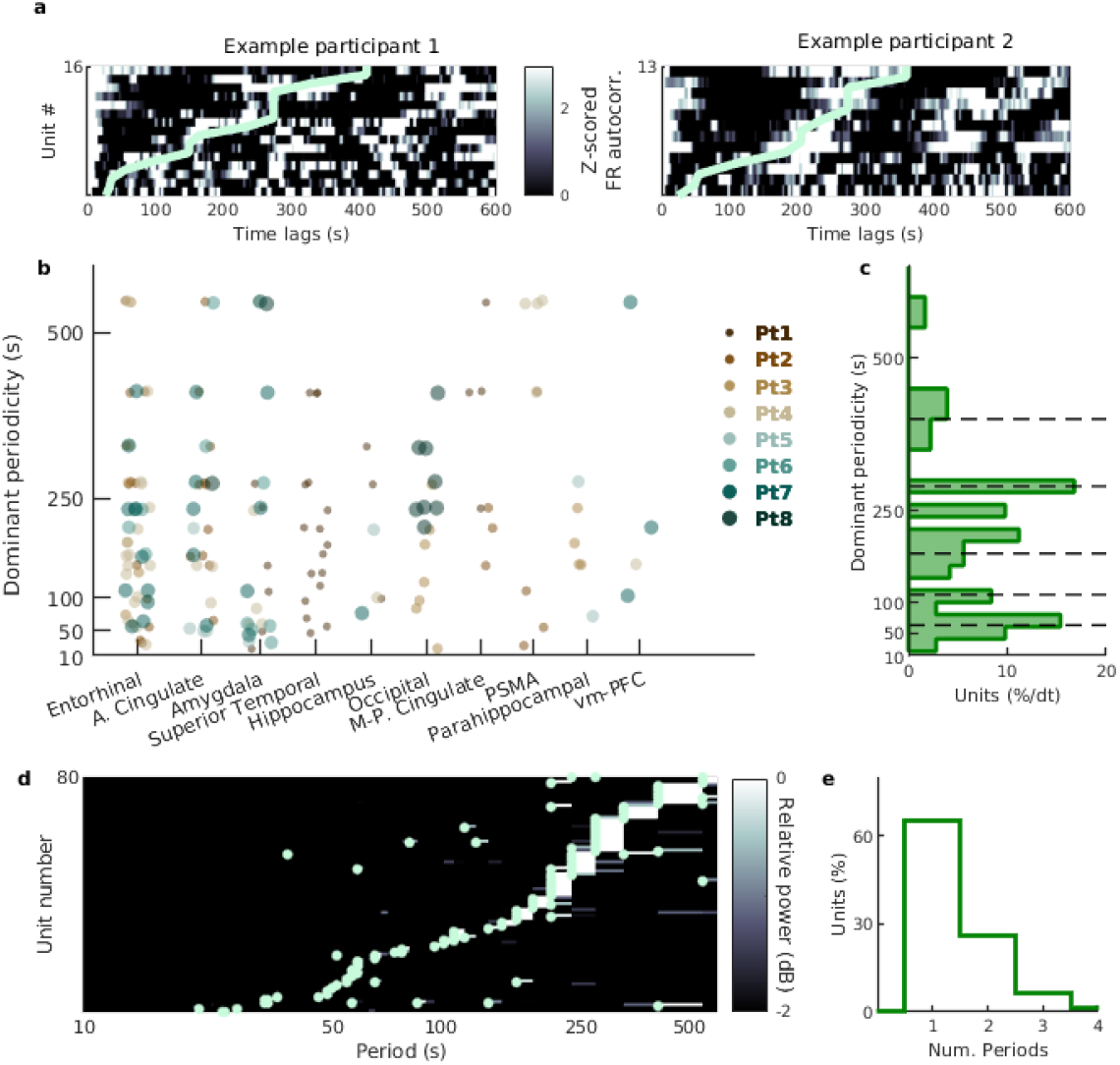
Distributions of the TPCs’ timescales. **a**. Distributions within subjects. Z-scored autocorrelation of the TPCs’ firing rates (colormap) for two example participants sorted by the dominant periodicity (light green line) for each unit (each row). **b**. Dominant periods of TPCs are shown within each region and for each participant (different colored/sized circles). **c**. The distribution of the dominant periodicity of all TPCs was not uniform (*p* < 10^−5^; single sample Kolmogorov-Smirnov test against uniform distribution). Due to the non-uniform bins, the percentage of units in each bin is normalized by the duration of the time bin. Note the pronounced peaks at 62.5s, 112.5s, 180s, 290s and 400s (marked with dashed lines). **d**. To determine prominent oscillations at periods other than the dominant periodicity, we examined relative power of the z-scored auto-correlogram (with respect to the maximum power at the dominant periodicity) for each unit (row) sorted by the dominant period. Light green circles indicate periods at which power was at least 75% of the maximum power (corresponding to the dominant period). **e**. Using the method in (d), we found the number of prominent periods (including the dominant period) for each unit. Shown is the distribution of the number of periods per unit and 35% of the units had prominent periodic activity in addition to their dominant periodicity.

### Time could be decoded from the population activity of TPCs

Given that TPCs exhibit periodicities at different timescales,, it should be possible to decode time from TPCs’ population activity–akin to a Fourier decomposition using periodic basis functions. To test this, we first partitioned the duration of the movie into equally sized epochs (bin durations for the epochs ranged from 1-90 seconds). We used linear discriminant analysis with a holdout approach to predict the time epoch within the movie using the firing rate of the TPCs as input features (Methods, Electrophysiological Analyses). We found that for bin durations longer than 6 seconds we were able to successfully decode time from the movie onset and the accuracy of the model, applied on an independent test set, was significantly above chance level (decoding time from shuffled TPCs’ firing rates; Fig. 4). The ability to extract precise, localized, temporal information from the population of TPCs, but not the shuffled data, shows that the periodic activity of the TPCs constitutes a viable mechanism to encode time. How the hippocampus may integrate such temporal information and incorporate it into encoding and retrieval of episodic memories deserves further investigations(*25–27*).

**Figure 4.**
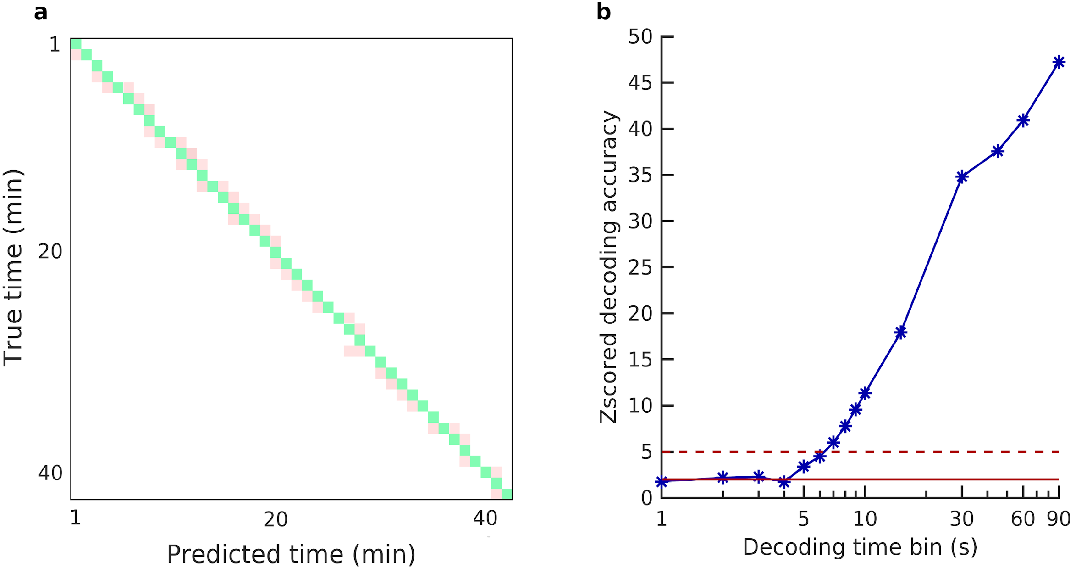
Decoding time from the population activity of the TPCs. **a**. Example confusion matrix (of the test set) from the time decoding analysis. Here, the time within the movie, and thus the activity of the TPCs (N=80), was divided into one-minute-long epochs and used as the input feature, while the output vector corresponded to the time bin numbers. Shown are the correctly classified time bins in green (the diagonal) and incorrectly classified time bins in pink (off diagonal). **b**. Decoding accuracy of the model on the test set was z-scored with respect to the shuffled data (decoding accuracy on shuffled TPCs’ firing rates) for different decoding bin sizes. For epochs larger than 6 seconds in duration, decoding accuracy was significantly above chance level (z=5; red dashed line).

### TPCs’ periodicities showed invariance with respect to narrative content

Can the presence of periodicity in the firing activity of the neurons be explained by the particular events and structure of this movie? First, we asked whether the cuts in the movie—defined as consecutive frames between sharp transitions(*20*)—were responsible for eliciting the TPCs’ periodic firing. However, the cut durations were markedly shorter (median, [25^th^, 75^th^] = 2.31s, [1.37, 3.10]s) than the TPCs’ dominant periodicities. Second, it appears unlikely that the TPCs’ timescales follow the content of the episode (e.g., the presence of specific characters in the movie was sparsely distributed; see Figure S6 in *ref*. 20). Further, the participants had not previously watched the episode and, therefore, could not predict the upcoming content that could, in return, dictate increase or decrease of firing activity. Lastly, if the TPCs’ periodicity was modulated by the content, one would expect that the activity of TPCs with similar dominant periodicities would be similar and, thus, highly correlated in time. This was not the case in our data and the distribution of correlation coefficients between adjacent TPCs’ firing (defined as TPCs with dominant periodicities within a certain time interval, e.g., 5s, 10s, 20s) was not significantly different from zero (*p*>0.05 for all intervals, signrank test). However, one cannot fully rule out the possibility that the neuronal firing was partly modulated by nested event boundaries of the narrative content(*28*).

To further assess the extent to which the TPCs’ periodic firing was modulated by external stimuli, we recorded data from additional five participants who watched the same episode but each half of the episode was presented to them at different speeds. In three of the participants, the first and second halves of the episode were played at regular and 1.5x speed respectively. In the other two participants, the order of the two speeds was reversed. Of the 285 recorded units (Table S1), we identified 80 units that exhibited TPC-like behavior during both conditions (regular and faster speeds) using the methods described earlier (Methods, Electrophysiological Analyses). Of the 53 units recorded from the entorhinal cortex, 19 (35.85%) were TPCs—a percentage similar to that observed in the nine participants described previously (34.43%).

If the periodicities of TPCs were merely determined by the content of the narrative, one would expect the periodicities to change in concert with the different rates of information in the two conditions. In contrast, several TPCs maintained the dominant periodicity of their firing rate during regular- and faster-speed movie viewing (Fig. 5a). These units exhibited stable periodic behavior across the two conditions (Fig. 5b), suggesting that their periodicity was independent of the narrative content. Overall, a significant fraction of the recorded TPCs (20 of 80 total; 25.00%, [15.99, 35.94]% binomial test confidence intervals) maintained their timescales between the two conditions (Fig. 5c).

**Figure 5:**
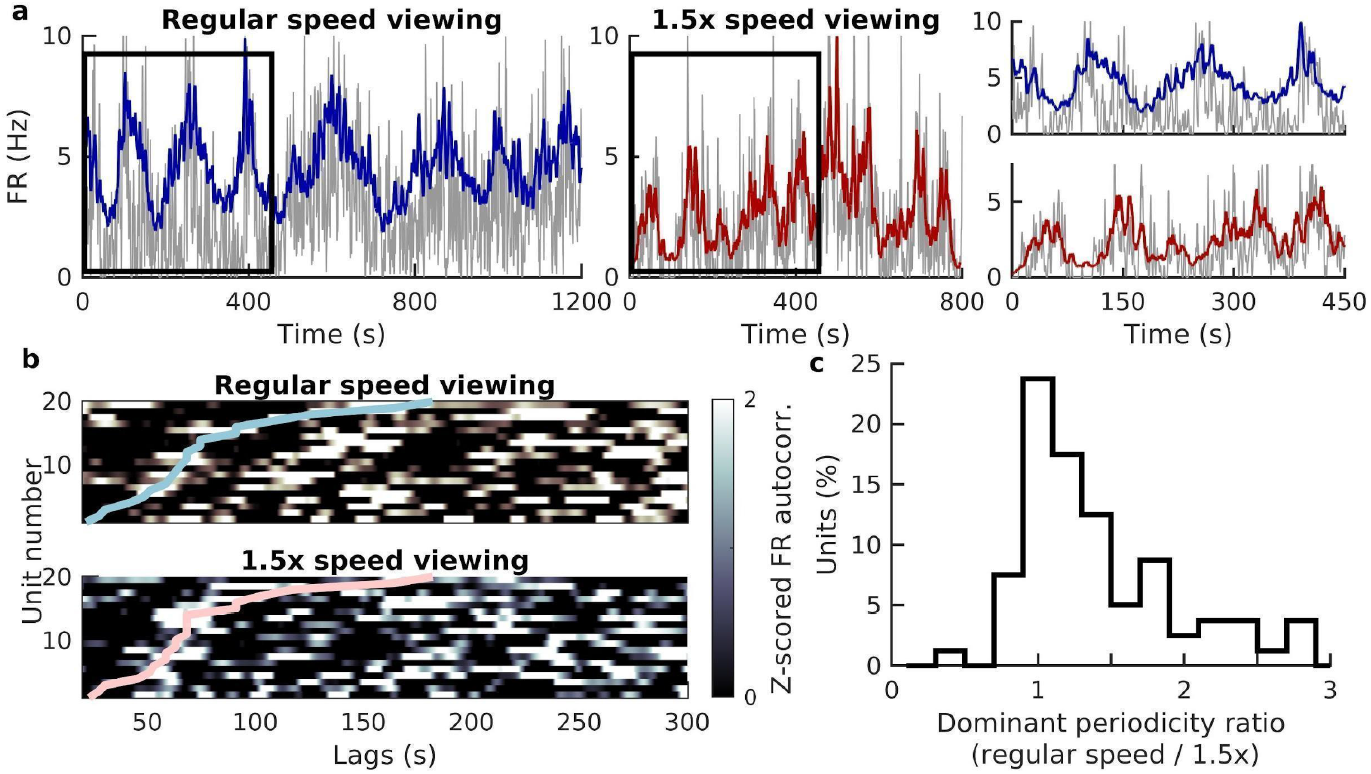
Maintained periodicity of TPCs during movie viewing at altered playback speeds. **a**. Left) Example unit’s firing rate (gray) during the first half of the episode played at regular speed overlaid with the GLM-fitted firing rate (blue). Middle) Firing rate of the same unit during the second half of the episode played at 1.5x speed overlaid with GLM-fitted rate (red). Right). Zoomed in views of the unit’s firing rate during the time intervals marked with black rectangles (Left and Middle). Note the same periodicity during movie viewing at regular speed (top) and accelerated speed (bottom). **b**. Z-scored firing rate autocorrelations of the units that exhibited the same periodicity during regular-speed movie viewing (top) and faster-speed movie viewing (bottom). Note that the neuron number is shared between the two panels and the colored lines represent the dominant periodicity of each unit. **c**. For each unit, the ratio of the dominant periodicity between regular-speed movie viewing and faster-speed movie viewing was computed. Shown is the distribution of this ratio across all identified TPCs (N=80). Of these TPCs, a significant percentage (25.00%, [15.99, 35.94]% binomial test) maintained their periodicity between the different speed conditions (defined as <10% change in their dominant periodicity across conditions).

### TPCs’ dominant periodicities remapped during memory test

Lastly, we asked whether the periodic activity related to the formation of episodic memories. We evaluated the periodic properties of TPCs during the memory test following the movie viewing (Fig. 1b). We employed the methodology described earlier to assess the significance of periodicity, as well as the dominant periods of the TPCs when participants were shown short clips and were tested for recognition memory (Methods, Behavioral Tasks). The majority (96.25%) of the TPCs maintained significant periodicity during the memory test, albeit at shorter timescales (Fig. 6a; Fig. S4). Although some units maintained their dominant periods during the memory test, most units (70.13%) “remapped” their periodicity to shorter timescales (Fig. 6b, c). The TPCs’ shorter periodicities during the memory test was not merely a response to the clip onsets as the time between clips (median, [25th, 75th] = 4.40, [3.38, 5.44] seconds; Fig. 6c, right) was much shorter than the dominant periods observed in the TPCs (Fig. 6c, left). Overall, the TPCs’ dominant periodicities were significantly shorter during the memory test compared to movie viewing, both on a population level (Fig. 6c, p=3.16×10^−5^, Wilcoxon ranksum test), as well as on the same cell basis (Fig. 6d; p=0.001, signrank test). It is worth noting, however, that although most units reduced their dominant periods during the memory test, few TPCs within the entorhinal cortex maintained or increased their dominant periods (27.59, [12.73, 47.24]%; binomial test; Fig. 6e). Whether the compression of the TPCs’ timescales during the memory test is relevant for individual behavioral performance and memory, remains to be explored in future investigations and will likely require technologies enabling sampling of a much larger number of neurons in humans.

**Figure 6:**
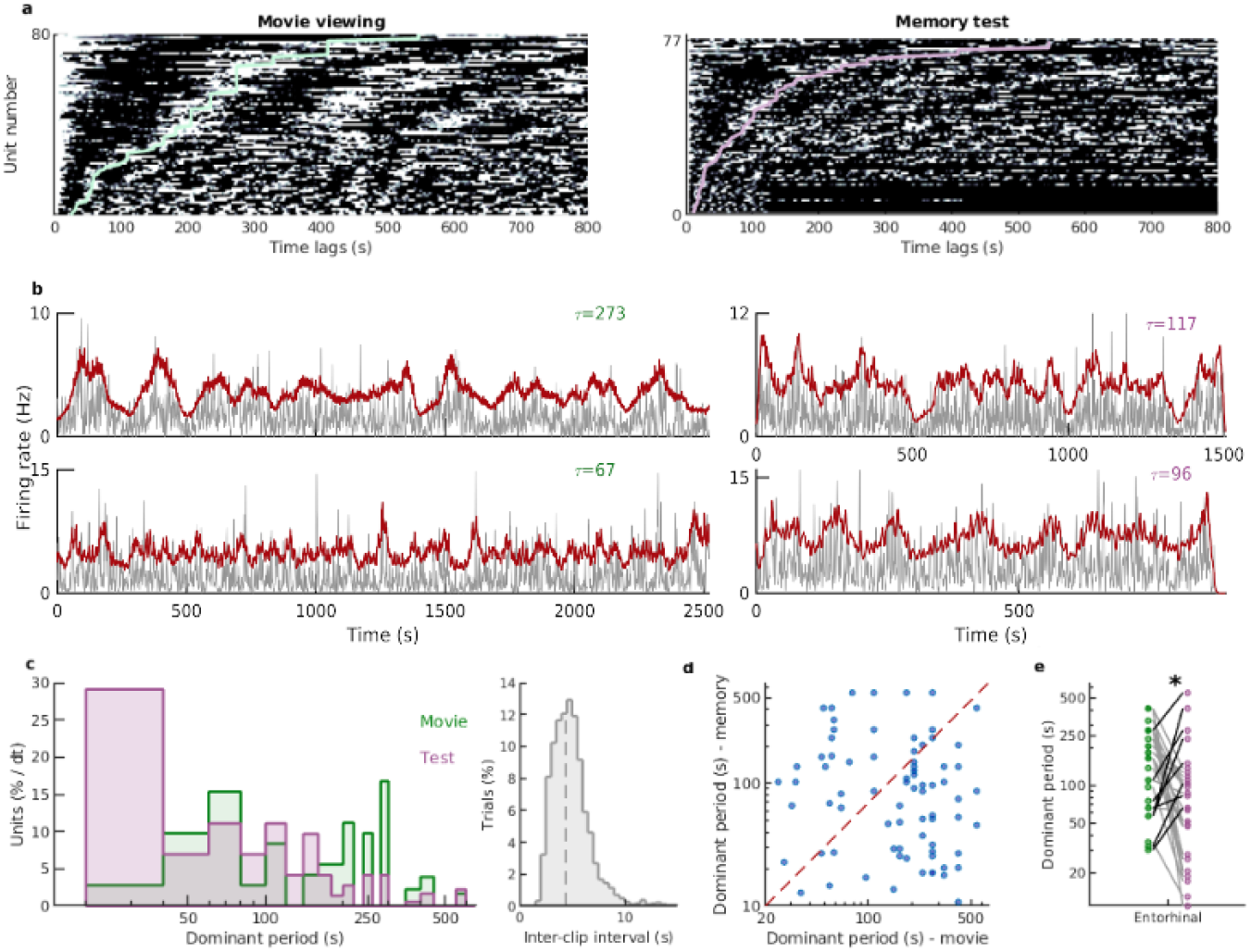
Periodic properties of TPCs during the memory test. **a**. Left) Z-scored autocorrelation of the TPCs’ firing rate (colormap) during movie viewing sorted by the dominant periodicity (light green line) for each unit (each row)(same as Fig. 2d reproduced here for comparison purposes). Right) Same as left but for the memory test. Of the 80 TPCs recorded during movie viewing, 77 (96.25%) remained as TPCs. **b**. Two example TPCs’ firing rate during movie viewing (left) and the memory test (right) recorded from the Entorhinal and cingulate cortex respectively. Gray line indicates the smoothed firing rate and the red line indicates the GLM-fitted firing rate. The value tau is the dominant period of the unit in each condition. **c**. Left) The dominant periods of the units were significantly shorter (*p*=3.16×10^−5^, Wilcoxon ranksum test) during memory test (n=77, purple distribution) compared to movie viewing (n=80, green histogram). Due to non-uniform time bins, the number of units per bin is normalized by the duration of the time bin. Right) The distribution of the inter-clip intervals during the memory test. Note that even the shortest dominant periods are longer than the inter-clip intervals shown here. **d**. For the same unit, the dominant period was shorter during the memory test compared to movie viewing (n=77, *p*= 0.001, signrank test). Red dashed line indicates the diagonal. **e**. For the TPCs recorded from the entorhinal cortex, shown are the dominant periods of the same cell during movie viewing (green circles) and memory test (purple circles). Gray (black) lines correspond to the units that decreased (increased or maintained) their dominant periods. A significant percentage of the TPCs within the entorhinal cortex maintained or increased their dominant periods during the memory test compared to movie viewing as indicated by asterisk (27.59, [12.73, 47.24]%; binomial test confidence intervals).

## Discussion

Recent studies in rodents have identified several cell types with time-dependent firing rates(*5–14*), notably hippocampal “time cells”(*6, 7*) and lateral entorhinal “ramping cells”(*14*). There have been similar quests in primate electrophysiology to discover neurons with time coding properties. The activity of temporal context cells in the monkey entorhinal cortex(*29*) aligns primarily with the rodent lateral entorhinal ramping cells. Recent human studies employing learning of sequences of word or picture stimuli described cells resembling the time and ramping cell types(*30–31*). It appears that time cells and ramping cells might contribute to two distinct types of temporal information: the sequential activity of time cells can map the delays with respect to a salient event along the time axis whereas the gradual change of activity of ramp cells in response to a salient event, which occurs at different time constants, may serve as a Laplace transformation of the elapsed time(*32*).

The time-dependent cellular machinery that we describe here is different altogether from those two cell types. It consists of a unique population of neurons with *periodic* modulation of activity across multiple timescales from tens of seconds to minutes. The reason that these cells so strikingly declared themselves is likely because of the continuous uninterrupted flow of information characterizing the current study. The key property of these cells was their periodicity over nearly an hour of relatively stable context yet with enormous variability in sensory input. This stability of temporal periodicity was further demonstrated by the fact that a subset of TPCs maintained their dominant periodicity despite the change in the video playback speed. This invariance to sensory input is required from an elementary neuronal clock where temporal information can be extracted from a population of neurons that together span a rich range of temporal scales from seconds to many minutes. In fact, previous models had proposed mechanisms that involved the extraction of time from a subset of neurons with periodic properties(*25, 26*).

Although the periodicity of the TPCs is observed in time and it is possible to decode time from the population activity of these cells, they may be responding to other time-varying signals rendering time representation a byproduct of this process. This argument may indeed hold true even for other types of time-coding cells and raises philosophical issues on whether time exists beyond “change” and the occurrence of events. Thus, perhaps the main significance of these findings is the presence of such temporal periodicity at the single neuron level at multiple timescales reaching many minutes and their primary presence in. the human entorhinal cortex.

The remapping of TPCs’ periodicities seen in the memory task following movie viewing may be related to multiple factors including memory, change in the temporal structure of the task, and change in context. It might also explain why such large-scale temporal periodicity has not been reported given that the recognition portion of the task more closely resembles the traditional stimulus-response task structure often employed in the field of human electrophysiology. If the shortening of periodicity is related to memory performance, these cells may play a role in temporal compression of experience required for memory retrieval(*32, 33*).

Of note, most of the entorhinal TPCs were in the anterior part of the entorhinal cortex. In humans, a recent fMRI study demonstrated that the activity of the anterolateral part of the entorhinal cortex is implicated in a temporal judgment memory task(*34*). Comparative anatomical studies of the human and rodent entorhinal cortex suggest that, in fact, the rodent LEC corresponds to the anterolateral portion of the entorhinal cortex and is, by nature, more multisensory compared to the MEC(*35*). Hence, it is possible that the TPCs might provide an additional temporal dimension to the incoming multisensory inputs to the entorhinal cortex.

The temporal periodicity of the TPCs begs comparison to *spatial* periodicity of grid cells. If a regular grid is a tessellation of n-dimensional Euclidean space, TPCs may be viewed then as one-dimensional *temporal* grid-like cells. Just like grid cells provide a multiscale map of a two-dimensional spatial environment, TPCs in humans may provide a multiscale map of the one-dimensional temporal environment. Akin to remapping of grid cells with change in size of the spatial environment(*15, 36*), TPCs exhibited remapping when the temporal structure of the task changed. These cells were by far most prevalent in the entorhinal cortex, but they were also found in ∼25% of anterior cingulate cells. Curiously, both entorhinal cortex and anterior cingulate were the brain regions where we had previously identified neurons with grid-like properties during human spatial navigation(*37*). Further, the entorhinal and anterior cingulate cortices were both implicated in retrospective duration estimations during encoding of long narratives(*38*).

It is possible that the periodic activity of TPCs may be related to the infra-slow (<0.1Hz) oscillations, previously described in the fMRI BOLD signals, LFPs, as well as single unit activity(*39–43*). The reported infra-slow activity was predominantly observed in sensory and association cortices, whereas the majority of the TPCs were recorded from the entorhinal cortex. It is possible that entorhinal cortex that receives convergent inputs from these areas(*35*) may integrate such infra-slow inputs into a more robust periodic time signal.

It should be borne in mind that there might be other interpretations for our findings. First, these TPCs were observed in epilepsy patients and, thus, it cannot be ruled out that periodicity is affected by epileptogenicity. However, the majority (95%) of the TPCs in the current study were recorded from regions outside the focus of seizure onset. Second, the periodic activity of the TPCs may subserve a range of behaviors, unrelated to time processing (e.g., chunking of experience at multiple timescales or efficient dynamics for neural communication). Lastly, it is likely that TPCs have conjunctive representations along dimensions other than time—a property that, if true, bears a resemblance to the conjunctive representation of navigational variables in the entorhinal grid cells(*44*). The potential synergy of grid cells and temporally periodic cells in providing spatiotemporal metrics of experience, and how their input may be incorporated in the hippocampus warrant further investigations, novel paradigms, and technological developments enabling concurrent recordings from large populations of cells in the human brain.

## Materials and Methods

### Participants

Participants were 14 epilepsy patients (age = 31±9; 9 Female), implanted with intracranial depth electrodes for seizure monitoring. Informed consent as obtained prior to the surgery and experiments were done in accordance with the Institutional Review Board at UCLA.

### Behavioral Tasks

The behavioral task (programmed in PsychToolbox, Matlab) consisted of participants watching an episode of the TV series 24 (season 6, episode 1, duration ∼ 42 minutes) on a laptop. Afterwards, they were presented with short clips (duration = 1.91 ± 0.72 s) and were asked to make a choice on whether they had seen the clip or not (response time duration = 2.39 ± 1.66 s), using the keyboard. The clips were divided into targets (clips chosen from episode 1 that they had just watched) and foils (clips chosen from episode 2 that they had never seen). The episodes of this series happen in consecutive hours of the day and, therefore, the characters’ appearances are very similar in the target and foil clips. Performance accuracy for each participant was computed as follows: (TP+TN)/(TP+TN+FP+FN), where TP, TN, FP, and FN are the true positive, true negative, false positive, and false negative respectively. We also computed an alternative behavioral performance measure, specifically d’ (d-prime) using the hit rate and false alarm rate values. These two measures of behavior (accuracy and d’) were highly correlated (*r*=0.974, *p*=4.20×10^−5^, Pearson correlation). The number of presented clips, and hence the duration of the memory test, varied from participant to participant.

Five additional participants performed an alternative version of the task. They watched the same episode of the TV series 24 but each half of the episode was presented at different playback speeds. In participants 1,3, and 5, the first half was presented at regular speed and the second half was presented at 1.5x speed. In participants 2 and 4, this order was reversed.

### Data Acquisition

Electrophysiological data was recorded from implanted electrodes that terminated in a set of nine 40 micro-m Platinum-Iridium microwires(*45, 46*). The number of electrode bundles, as well as their locations, were different for each participant and determined solely by clinical criteria. Wide-band local field potentials were recorded from eight microwires (the 9th microwire was used for referencing) using a 128-channel (or 256-channel) Neuroport recording system (Blackrock Microsystems, Utah, USA) sampled at 30 kHz.

### Electrode Localization

A high-resolution post-operative CT image was obtained and co-registered to a pre-operative whole brain and high-resolution MRI for each participant using previous methods (Fig. 1c; Table S1). The locations of the microelectrodes were determined by examining the location of the electrode artifact on the co-registered images. For further details, see ref. *21*.

### Electrophysiological Analyses

Data were analyzed offline using custom code as well as functions and toolboxes in Matlab and Python.

#### a) Spike detection and sorting

Spike detection and sorting was done using previous methods(*21–24*). Briefly, we applied a bandpass filter to the broadband data in the 300-3000Hz to detect spikes that were subsequently sorted using the Wave_clus toolbox. Furthermore, the automatically-detected clusters were manually inspected for: 1) spike waveforms; 2) presence of refractory spikes; as well as 3) the ISI distribution for each cluster. Clusters with firing rates below 0.05 Hz were discarded from further analysis. Note that the movie viewing and recognition memory test phases were recorded within a single session and, thus, spike detection and sorting was performed over the entire session. The activity of each unit was then separated for each phase (viewing/memory) of the experiment.

#### b) Firing rates and their autocorrelations

A time vector with a bin size of 100ms was constructed and, for each unit, the number of spikes within each time bin was computed. This raw spike train was used for the GLM analyses (next section). The smoothed spike trains were computed using a 0.5s Gaussian smoothing kernel on the raw spike histograms, which were then converted to firing rates after division by the duration of the time bin (Fig. 2a, 5a, 6b, S2, S3, S4). To inspect the presence of putative oscillations in the spiking activity, a normalized autocorrelation was computed over the smoothed firing rate.

#### c) Determining significant temporally periodic cells (TPCs)

To determine whether the periodicity in the spiking activity, as demonstrated by the autocorrelation of the firing rates, was statistically significant, we used a shuffling procedure. For each unit: 1) we chunked the firing rate into 1-second-long segments and randomly shuffled the segments in time (x 250); 2) the previous step was repeated for 2-second-long segments. This procedure yielded 500 shuffled firing rates for which an autocorrelogram was calculated. Next, we compared the autocorrelation of the true firing rate against the autocorrelation of the shuffled firing rates. Units with true autocorrelations that had values beyond the 2.5% and 97.5% of the shuffled data were identified. Further, we used a cluster-based permutation test(*47*) to correct for multiple comparisons (given the large number of lags that were being tested). Specifically, we used the function permutation_cluster_test from MNE Python package(*48*) and units with significant clusters were deemed to be TPCs. The different steps of this procedure are demonstrated in Fig. S2.

#### d) Generalized Linear Models (GLMs)

The time-varying firing rate of each unit was modeled as an inhomogeneous Poisson process(*49*) using basis functions that are periodic in time:

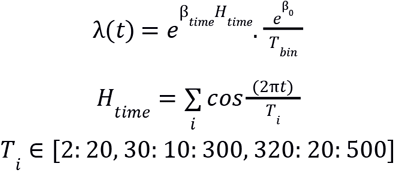

Here, T_bin_ is the bin size in time (0.1 s), H refers to the design matrix associated with the temporal covariates, in this case cosine functions with different periods (T_i_), and betas are the parameters associated with the design matrix in time and a constant term. Note that the exponentiation is done element wise in this case. This allowed us to determine the periods (T_i_) that significantly contributed to the firing activity of the units (p<0.001). Oftentimes, units had more than one significant term. The distribution of these periods is shown in Fig. S1.

#### e) Dominant periodicity

To determine the strongest oscillation periodicity in the firing rate of the TPCs, we z-scored the autocorrelation of the smoothed firing rates (described in b) with respect to the shuffled data (described in c), referred to as z-scored autocorrelation for simplicity (Fig. 2d). Next, we performed FFT analysis on the z-scored autocorrelation values for each unit and the period with the maximum power was chosen as the dominant period of the unit (Fig. S2). To assess the strength of other potential periodicities, the power was normalized with respect to the strongest peak (corresponding to the dominant periodicity) and peaks with 75% of the maximum power were considered as secondary, tertiary, etc. periodicities (Fig. 3).

#### f) Decoding time from TPCs’ population activity

Decoding analysis was done using Linear Discriminant Analysis as a classification method. We divided the data into equally sized time epochs and we performed this analysis for different bin sizes of [1:10, 15, 30, 45, 60, 90] seconds. The epoch number was used as the output of the classification model and the activity of the TPCs within each epoch was used as the input to the model. Further, we used a hold-out method, i.e., the model was trained on randomized 75% of the data and an independent 25% of the data were left aside for testing and the model performance was evaluated on the test dataset (Fig. 4). Additionally, the performance of the model was compared against shuffled data: the same classification method was applied on the temporally shuffled activity of the TPCs. For each unit, we chunked the firing rate into 1-second-long segments and randomly shuffled them in time. We then concatenated the shuffled firing rates of all TPCs and obtained a surrogate input. We applied the same classification method on the shuffle data and computed model accuracy. We repeated this shuffling procedure 250 times.

## General

We thank Matias Ison, Hanlin Tang, Jie Zheng, and Guldamla Kalender for assistance with data collection and processing. We also thank the participants for taking part in our study.

## Funding

This work was supported by the NIH NINDS (NS033221 and NS084017 to I.F.) and NSF Center for Brains, Minds and Machines and McKnight Foundation (G.K.)

## Author Contributions

Z.M.A., G.K., and I.F. conceived and designed the research. Z.M.A. and I.F. performed analyses. I.F. performed all implantation of the depth electrodes. Z.M.A. and I.F. wrote the paper. All the authors commented on the manuscript.

## Competing interests

Authors declare no competing interests.

## Data and materials availability

Custom code used and datasets analyzed for the current study are available from the corresponding author upon reasonable request.

## Figures and Tables

For ease of readability, we have included the main figures and legends within the text.

## Supplementary Materials

**Fig. S1.**
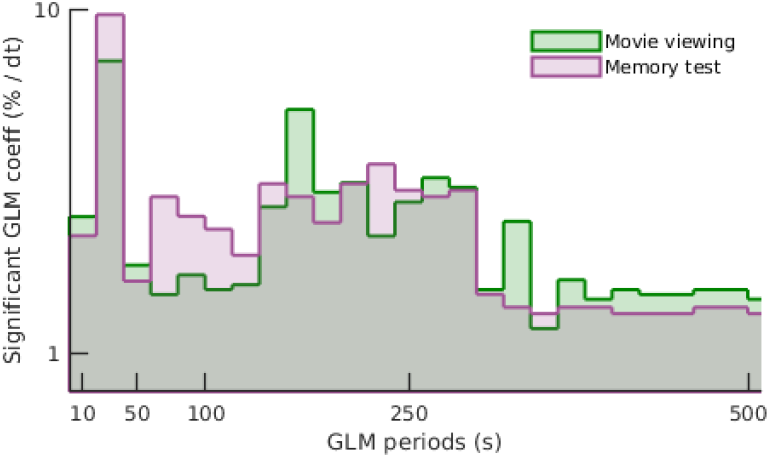
Significant periods in the GLM-fitted firing rates. The distribution of the periods that were deemed significant (p<0.001) in describing the time-varying firing rate of the units during movie viewing (green; number of significant periods detected by the GLM method = 787 from 80 units) and memory test (purple; N = 589 from 77 units). For the majority of the units, the GLM fitting resulted in more than one term that contributed to the firing. Note the shift towards shorter periodicities during the memory test compared to the movie viewing (*p*=0.01; Wilcoxon ranksum test).

**Fig. S2.**
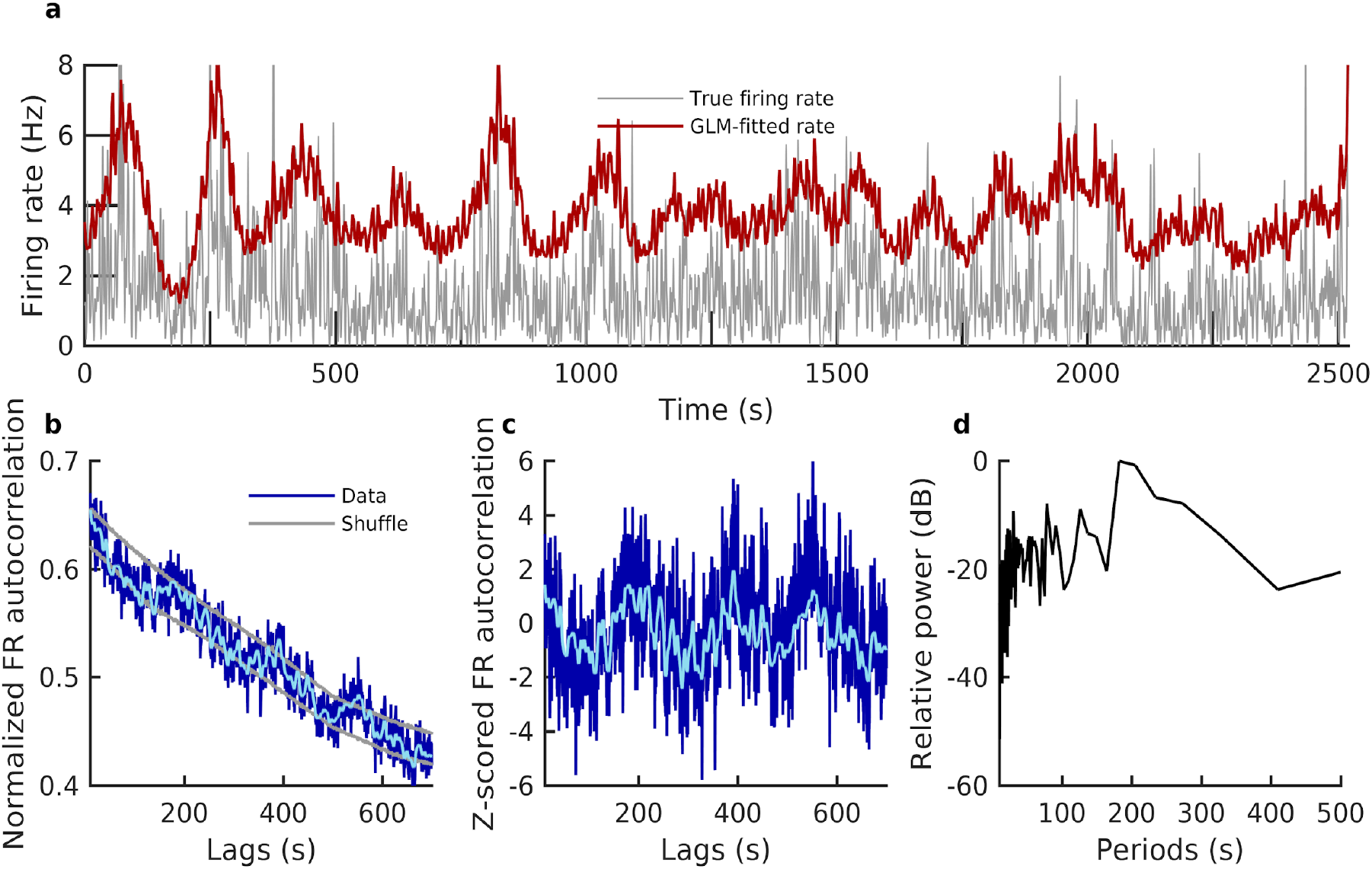
Steps to determine significant TPCs and their dominant periods. **a**. Example TPC from the left entorhinal cortex; smoothed firing rate (gray) overlaid with the GLM-fitted rate (red). **b**. Normalized autocorrelation of the smoothed firing rate shown in (a) is shown in dark blue. Gray lines indicate the 2.5% and 97.5% of the shuffle data (see Methods, Electrophysiological Analyses), which were obtained from generating shuffled firing rates, followed by computing the autocorrelograms of the shuffled rates. A unit with firing rate autocorrelations beyond the shuffled data was then corrected for multiple comparisons (using cluster-based permutation test), and units with significant clusters were deemed to be TPCs. **c**. The true firing rate autocorrelogram was z-scored with respect to the shuffled data (subtracting the mean and division by the standard deviation) and is shown in dark blue (light blue curve is the smoothed version of the z-scored autocorrelation and is shown only for visualization purposes). **d**. Relative power (FFT normalized by the maximum power) of the z-scored firing rate autocorrelogram (c), which was used to compute the dominant periodicity of the units’ firing rates. Dominant periodicity was defined as the period that contained the maximum power (∼200s for this unit).

**Fig. S3.**
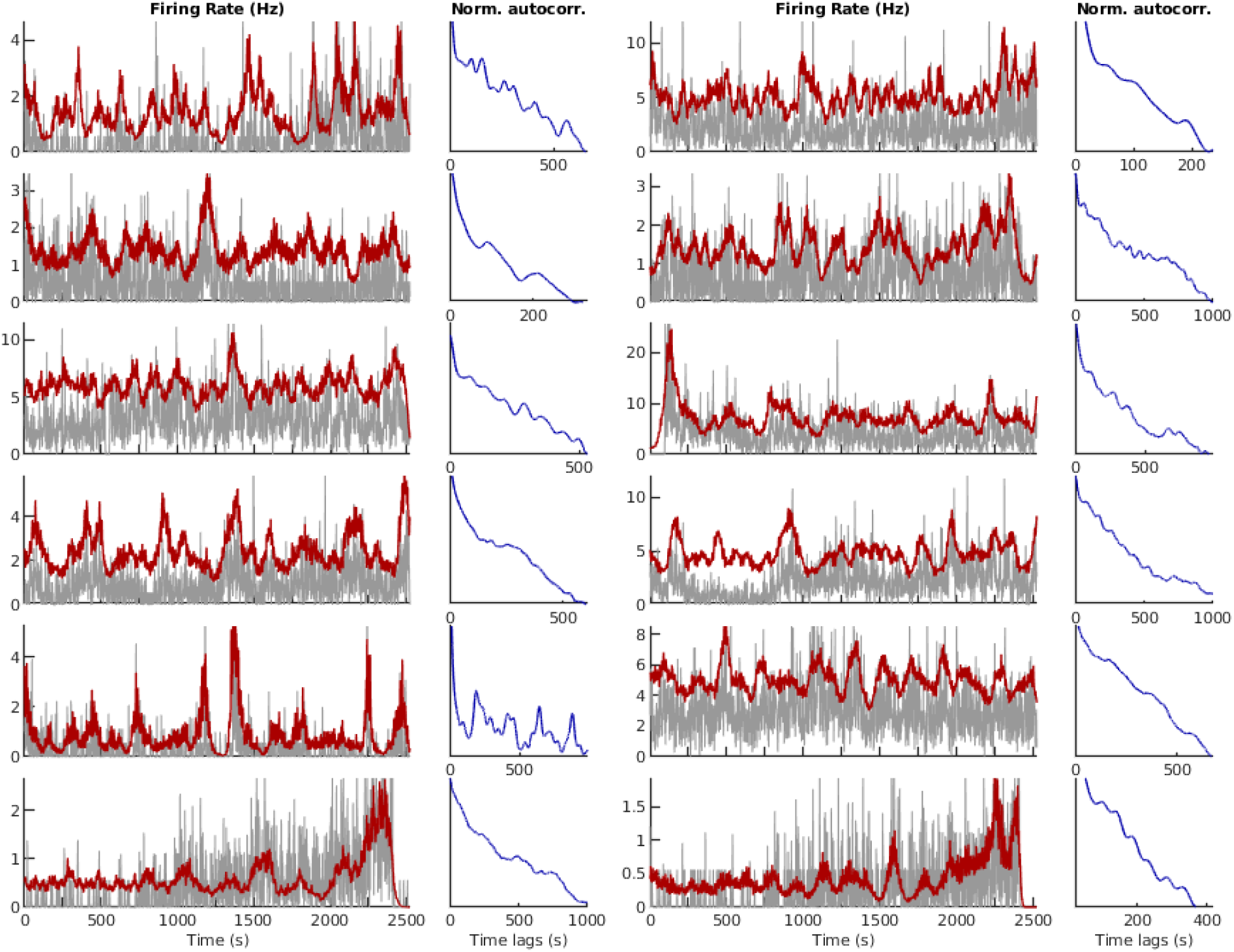
Example TPCs during movie viewing. Additional examples of 12 units that were identified as TPCs. Within each column: Left) Smoothed firing rate (gray) overlaid with GLM-fitted rate (red); Right) Normalized autocorrelation of the smoothed firing rate to demonstrate periodicity. These units were recorded from the following regions (from left to right within the rows): Mid. Post. Cingulate; Entorhinal; Entorhinal; Amygdala; Entorhinal; vm-PFC; Entorhinal; Entorhinal; Occipital: Entorhinal; Entorhinal; Entorhinal.

**Fig. S4.**
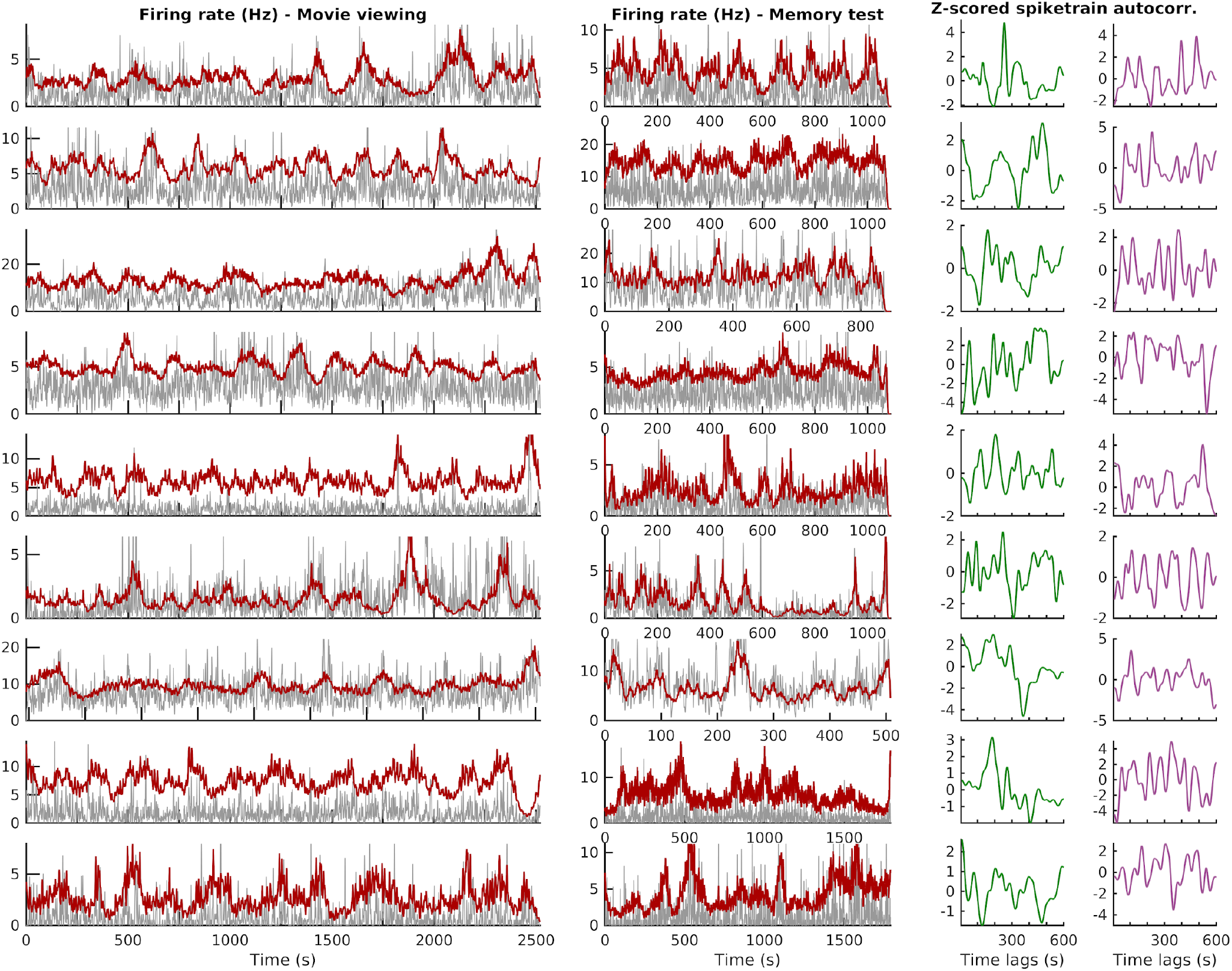
Example TPCs during movie viewing and their respective activity during the memory task. We tracked the activity of the same unit during movie viewing and memory test. Shown are nine different units (each row). Left) Firing rate (gray) and GLM-fitted rate (red) during movie viewing. Middle) Same as in (left) but during the memory test. Right) Z-scored autocorrelograms of the spike trains (smoothed only for visualization purposes) during movie viewing (green) and memory test (purple). These units were recorded from the following regions (top to bottom): Entorhinal; Entorhinal; Superior Temporal; Entorhinal; Amygdala; A. Cingulate; Entorhinal; Entorhinal; Entorhinal. Note the higher frequency oscillations (faster periodicity) during memory test (purple).

**Table S1.** Electrode localizations. For each participant (rows), all of the recording electrode locations that had units are listed (columns: categories that were used for group analysis). The hemispheric locations of the electrodes are marked with R and L, referring to the right and left hemispheres respectively. The numbers in parentheses indicate the number of electrodes within each hemisphere (if there were more than one). ^a^Additional units were recorded in the fusiform gyrus. Participants 10-14 performed an alternative version of the task that consisted of watching one half of the episode at regular speed and the other half at a faster speed (1.5x speed; see Methods).

| **Locs**<br>**Pt. ID** | <u>Entorhinal</u><br><u>Cortex</u> | <u>Hippocampus</u> | <u>Parahippocampal</u><br><u>Gyrus</u> | <u>Subiculum</u> | <u>Amygdala</u> | <u>Superior</u><br><u>Temporal</u><br><u>Gyrus</u> | <u>Anterior</u><br><u>Cingulate</u> | <u>Middle &</u><br><u>Posterior</u><br><u>Cingulate</u> | <u>Ventromedial</u><br><u>Prefrontal</u><br><u>Cortex</u> | <u>PreSMA</u> | <u>Occipital</u> | <u>Insula</u> |
| --- | --- | --- | --- | --- | --- | --- | --- | --- | --- | --- | --- | --- |
| **1** | - | L<br>R | - | - | R | L<br>R (2) | - | L<br>R (3) | - | - | - | - |
| **2** | L<br>R | - |  |  | - | - | R (2) | R |  | L<br>R | - | - |
| **3\*** | R | - | L<br>R | L | - | - | - | - | - | - | L (2) | - |
| **4** | L<br>R (2) | L | - | - | L<br>R | - | L<br>R | - | L<br>R | L<br>R | - | - |
| **5** | L | - | - | L | L | - | - | - | L | - | - | - |
| **6** |  | L | R | - | L<br>R | - | R | - | - | - | - | - |
| **7** | L | - | - | - | L<br>R | - | L | - | - | R | - | - |
| **8** | L<br>R | L<br>R | - | - | L | - | L<br>R | - | L<br>R | - | - | - |
| **9** | - | - | - | - | R | - | R | - | - | - | R(2) | - |
| **10<sup>a</sup>** | - | R | - | L | L | L | L | - | R<br>L | R | L | L |
| **11** | R | R | - | - | L | - | - | - | - | - | - | R (2)<br>L |
| **12** | R<br>L | R<br>L | - | - | L | L | - | - | L | - | - | - |
| **13** | L | R<br>L | R | - | L | R | - | - | R | - | - | - |
| **14** | R<br>L | R<br>L | - | - | L | L | R<br>L | - | R | - | - | - |

**Table S2.** Number of total units and significant TPCs recorded per region. The numbers reported here are from the original 9 participants.

| **Participant** | **Number of Units** | **Number of TPCs** |
| --- | --- | --- |
| *Hippocampus* | 19 | 4 |
| *Entorhinal Cortex* | 80 | 30 |
| *Parahippocampal* | 12 | 2 |
| *Amygdala* | 74 | 12 |
| *Superior Temporal* | 28 | 7 |
| *Anterior Cingulate* | 51 | 13 |
| *Middle & Posterior Cingulate* | 25 | 3 |
| *VM Prefrontal Cortex* | 21 | 2 |
| *PreSMA* | 20 | 3 |
| *Occipital* | 43 | 4 |

**Table S3.** Number of TPCs recorded per participant and their timescales. The numbers reported here are from the original 9 participants. TPCs’ timescales are represented by their dominant periodicities (shown are mean ± std within each participant).

| Participant | Number of TPCs | TPCs' timescales (s)<br>mean dominant period ± std |
| --- | --- | --- |
| 1 | 13 | 207.74 ± 125.12 |
| 2 | 16 | 198.74 ± 129.23 |
| 3 | 5 | 189.14 ± 74.68 |
| 4 | 15 | 172.54 ± 122.84 |
| 5 | 0 | NA |
| 6 | 4 | 126.86 ± 90.00 |
| 7 | 7 | 213.45 ± 187.23 |
| 8 | 15 | 198.50 ± 143.39 |
| 9 | 5 | 309.36 ± 140.24 |

## Notes

### Competing Interest Statement

The authors have declared no competing interest.

### Summary of Updates

Title change, as well as renaming units from periodic time cells to temporally periodic cells.

